# Unsupervised decomposition of massively parallel TraDIS-Xpress fitness data from *Escherichia coli* identifies distinct functional and expression modules of antimicrobial resistance determinants

**DOI:** 10.64898/2026.10.11.758300

**Authors:** Muhammad Yasir, A. Keith Turner, Andrea Telatin, Sarah Bastkowski, Emma R. Holden, Ian G. Charles, Mark A. Webber

## Abstract

Linking genotype to phenotype is a central goal in biology, one way to do this is using transposon insertion sequencing (TIS) where the fitness impacts of mutations across a genome are assessed in different conditions. Traditional analyses use comparison of each gene response independently in control versus test conditions, so a compendium of hundreds of test conditions returns hundreds of separate ranked gene lists.

To unlock the potential information hidden within large data sets, we applied independent component (ICA) analysis to 754 *Escherichia coli* BW25113 TraDIS-Xpress samples. These samples included a series of antibiotic and other stress condition exposures. ICA is designed to identify hidden structure within complex data and to extract meaningfully linked components. In this analysis each annotated gene was resolved into a gene body and 198 bp 5′ and 3′ features so that expression effects driven by the transposon’s outwards facing promoter enter the matrix separately from gene disruption effects.

The decomposition returned 99 components, which we term fxModules, which were defined as containing elements with similar responses across the whole compendium rather than per any individual experiment. Of the 754 stress samples, the antibiotic arm included 110 samples derived from exposure to 14 separate agents at multiple concentrations. Many of these have been highly studied allowing us to explore the accuracy of predicted responsive fxModules. This analysis showed excellent predictive power of the approach and the fxModules responsive to individual drugs typically reproduced known mechanisms at a resolution superior to analysis of single gene sets. For example, exposure to fosfomycin identified responsive fxModules containing genes which are known to be involved in uptake of the drug, cAMP-CRP control and exposure to gentamicin identified oxidative phosphorylation split across three fxModules responding with different magnitudes and directions at different concentrations. This recapitulates recent work studying the landscape of *E. coli* genes sensitive to this drug with high precision without any prior knowledge of functional annotation. Similarly precise predictions were made for other drugs.

Responsive modules also identified novel groups of genes and linkages beyond the boundaries of defined pathways as being responsive to various stresses demonstrating the ability to predict new genotype-phenotype links.

Inclusion of the regulatory regions also resolved fxModules which were entirely contributed by promoter regions, for example for chloramphenicol genes responsible for efflux were identified by signals in intergenic control regions, and unidirectional insertions in gene bodies, a signal that a gene level matrix would not be able to predict with accuracy.

These data show that ICA decomposition provides an efficient and accurate annotation independent approach to identifying sets of genes with related functional impacts in any given condition in the compendium. The functional modular association of genes also help in predicting gene functions of unknown genes based on the group function. This approach allows rapid identification of novel genotype-phenotype links and unlocks the power of multiple comparisons of transposon mutagenesis data.

## 1. Introduction

Transposon insertion sequencing (TIS) links genotype to phenotype at genome scale by measuring how insertion abundance at every locus change under selection (Langridge et al. 2009; Cain et al. 2020). This has been hugely important in understanding roles for genes in many conditions, for example in antimicrobial research where in a single, parallel experiment roles for genes involved in drug entry, target engagement, efflux and physiological states relevant to survival can be predicted (Yasir et al. 2020; Turner, Eckert, et al. 2020). Whilst TIS provides a wealth of information the standard output is one fitness statistic per gene per condition by comparison of mutant abundance between test and control conditions.

As costs have reduced, the throughput of TIS experiments has risen and single mutant libraries can now be assayed against dozens of conditions in parallel. However, the analytical approaches involved have largely not changed and each condition is typically still analysed in isolation, and each gene is still tested independently (Barquist et al. 2016; Page et al. 2020). Therefore, a compendium of hundreds of conditions returns hundreds of separate ranked lists whose relation to one another must be established afterwards by subsequent analysis of these outputs. Increasing the size of TIS datasets has therefore increased the number of separate results rather than the resolution of any one of them, and what is missing is a common set of coordinates in which every condition in a compendium can be described and compared directly.

Two further constraints of the per gene TIS output approach also need addressing. The first relates to the coverage of the genome. Conventional TIS cannot assay essential genes, which do not tolerate insertion, and it treats intergenic regions as uninformative. Versions of TIS such as TraDIS-Xpress address this by incorporating outward transcribing promoters within the transposons, so that insertions upstream of a gene drive its overexpression and insertions oriented into its 3′ end suppress it through antisense transcription (Yasir et al. 2020). Genes are therefore assayed by altered expression as well as by inactivation, and essential genes can become accessible for analysis, but the comparison analysis approach can make false positive calls due to small loci size of 5′ and 3′ regions.

The second constraint of current list based analysis approaches is each gene is tested independently, and any coordinated response across conditions is reconstructed after analysis by enrichment against pathway or regulon databases. Those databases were assembled for other purposes, and their boundaries need not correspond to the units that share a fitness consequence. A functional unit may split across several database categories, and a database category containing components with different requirements returns a single statistic that averages over them.

The approach of unsupervised matrix decomposition addresses the constraints of functional units and can be applied across a compendium of data rather than a single experiment. For example, independent component analysis (ICA) applied to bacterial transcriptomes has been used to recover independently regulated modules, or ‘iModulons’, whose membership is set by covariation in the data rather than by prior annotation and which correspond closely to known regulons(Sastry et al. 2019; Rychel et al. 2021). The same approach has also been applied to randomly barcoded Tn-seq fitness data in *Pseudomonas putida* to recover fitness modules, or ‘fModules’ (Borchert et al. 2024). Neither has been applied to fitness data in which expression effects are resolved alongside disruption effects. Here we extended this approach to a compendium of TraDIS-*Xpress* data. We recovered ‘fxModules’ for components defined jointly by fitness and expression, distinguishing them from iModules, which group genes by shared transcription regulation, and fModules, which group genes by shared fitness. This distinction is not cosmetic: the threshold selection procedure developed for either predecessor is not transferable to this data type, and the internal architecture of the resulting modules is different.

We analysed a compendium of 754 *Escherichia coli* BW25113 TraDIS-Xpress experiments at sub-gene resolution where promoter, repressor and gene body signals all entered the matrix as separate features and all conditions share one set of component definitions. We established that the membership thresholding procedure developed for iModulons and fModules did not transfer to this data type and determined a working cutoff for module assembly empirically. We characterised the internal architecture of the modules, which differs from that reported for either predecessor. We then used the antibiotic arm of the compendium (consisting of a subset of 110 conditions representing data from 14 agents assayed at four concentrations), as the test case. Known mechanisms of action and principal resistance determinants were effectively and independently established for most of these antibiotics using this approach in a single analysis. This provided improvements in speed with the single, parallel analysis providing the module memberships and precision of assignment of genes to phenotype compared with the individual exposure analyses previously completed (Bastkowski et al. 2026).

This approach shows that decomposition can effectively identify and predict fxModules which represent meaningful sets of genes which are involved in stress responses from compendiums of TIS data. The resolution is superior to current common analysis approaches, is not reliant on annotation and links groups of genes into functionally interacting pathways. Recall of known mechanisms is strong and predictions of new genes involved in a given stress condition can be made with confidence. We believe this method will be useful in unlocking the hidden potential available from large TIS datasets and can be a significant tool in the goal of linking genotype to phenotype.

## 2. Results

### 2.1 TraDIS-Xpress fitness and expression compendium

We assembled a compendium of 754 samples of *Escherichia coli* BW25113 TraDIS-Xpress sequencing data of antibiotic exposure, a variety of physical and chemical growth stress conditions and library controls (Yasir et al. 2020; Yasir et al. 2025; Yasir et al. 2022; Holden et al. 2022; Holden et al. 2021; Thomson et al. 2022; Turner et al. 2021; Turner, Yasir, et al. 2020; Bastkowski et al. 2026; Holden et al. 2025). All the samples are from the same high density mini-Tn5 mutant library of approximately 800,000 insertion mutants, with one insertion on average every 6 bp across the genome and were processed through a common mapping pipeline with insertion counts determined per annotation across the genome.

Unlike conventional transposon insertion sequencing, the TraDIS-Xpress transposon carries an outward transcribing, IPTG inducible tac promoter 3′ of the kanamycin resistance cassette. Insertions into the 5′ of a gene therefore drive its overexpression, while insertions oriented into the 3′ end suppress expression through antisense transcription, allowing genes to be assayed by altered expression as well as by inactivation. This includes essential genes, which cannot be assayed by insertional inactivation at all. To preserve this information, each gene was resolved into three separate features: the gene body, the 5′ region (198 bp) and the 3′ region (198 bp) (Figure 1A). Any 5′ or 3′ region overlapping more than 25 bp onto neighbouring genes was dropped from analysis. The decomposition was therefore performed on a feature matrix of 6,311 features (Figure 1B) instead of approximately 4,500 genes. Of these features, 4,556 (72%) are gene-body features (4,227 locus tags and 329 base coordinates), 1,219 (19.3%) are 5′ features and 536 (8.5%) are 3′ features.

**Figure 1:**
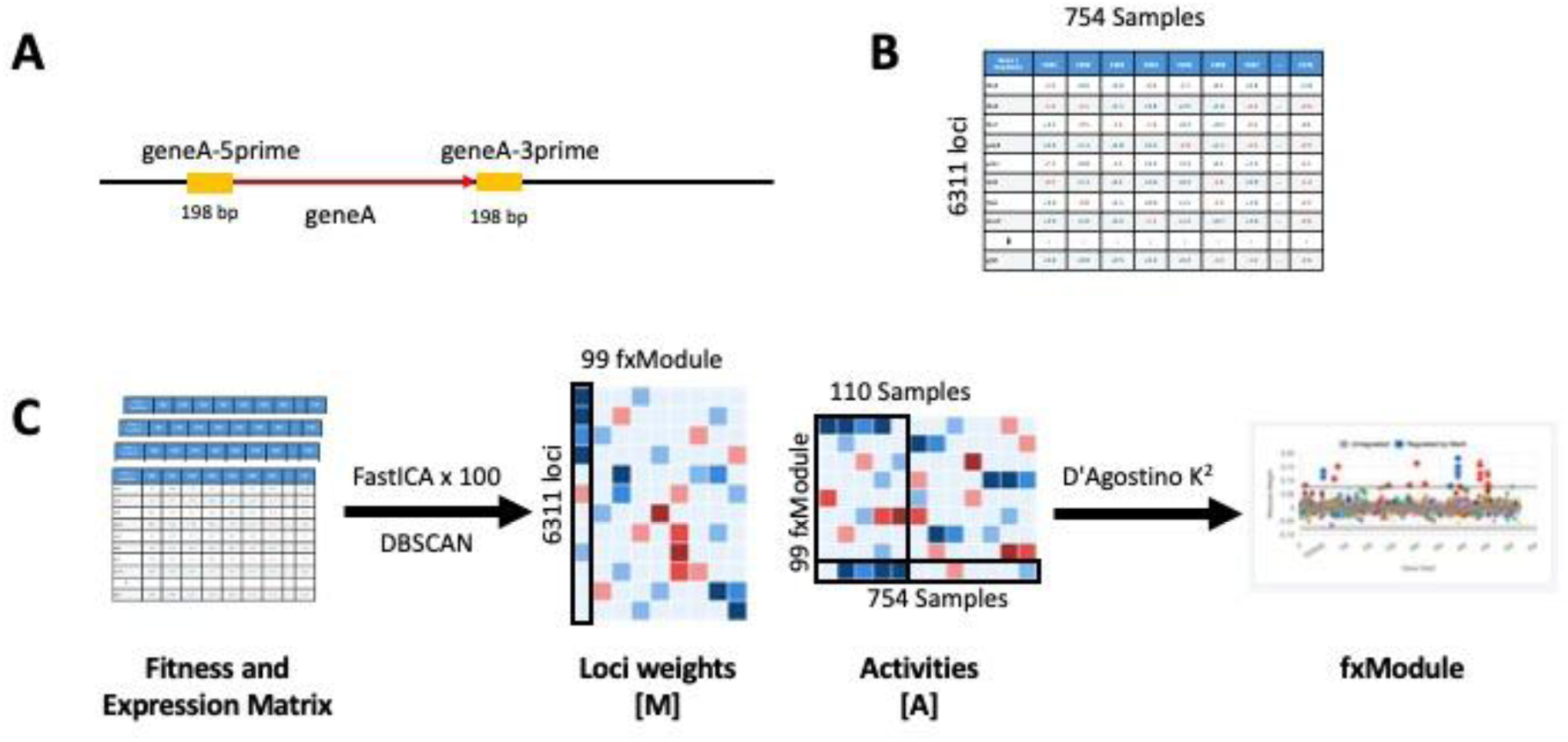
Sub-gene feature decomposition and ICA workflow. **(A)** Each gene is resolved into three features: the gene body and 198 bp 5′ and 3′ flanking regions. Flanking regions overlapping a neighbouring gene by more than 25 bp were removed, giving 6,311 features. **(B)** Input matrix of fitness and expression values for 6,311 features across 754 TraDIS-Xpress libraries. **(C)** FastICA is run 100 times on the full compendium with components clustered across runs by DBSCAN, giving a feature weight matrix M and a sample activity matrix A. One degenerate component holding the majority of features at every cutoff tested was removed as a global variance sink, leaving 99 fxModules. Membership is then assigned by the D’Agostino K² procedure on the source weights to define the fxModules.

ICA was applied to the full 754 samples compendium and not to any single experiment, so that component definitions are set by variation across the entire experimental space and module activities in any one experiment can be read against that common base. Downstream biological analysis focuses on the antibiotic exposure arm of the compendium, comprising 110 samples covering 14 antimicrobials (Figure 1C): the 13 agents reported previously by Bastkowski et al. (2026) together with chloramphenicol, each assayed at four concentrations relative to its minimum inhibitory concentration against *E. coli* BW25113, namely 0.25×, 0.5×, 1× and 2× MIC, in duplicate.

A module is defined by ICA with FastICA, which decomposes the features by sample fitness and expression matrix into components based on the dimensions according to the diversity in the samples, and in this case, it examined the data at 400 dimensions to resolve 99 modules (Supplementary Figure S1). This step assigns a weight to every feature (M-Matrix) and an activity to every sample (A-Matrix) (Figure 1C). Assigning features (members) to modules requires a threshold, D, on the ICA source weights, applied through the established D’Agostino K² procedure with a D-1350 to assign membership for this data set (Supplementary Figure S2).

### 2.2 fxModule activity across the antibiotic arm

The ICA centres each component across the whole 754 samples compendium, using this annotation independent module membership assignment approach, we asked whether module activities carry interpretable structure under a defined selection. Specifically, we explored whether antimicrobials differ in how far their fitness response is concentrated in dedicated components rather than distributed across shared physiology. From the compendium, a subset of 110 antibiotic stress samples is discussed in this manuscript. For membership of the modules, D’Agostino K^2^ value of D = 1,350, resulted in 51 modules with 3 or more than 3 members, with a median size of 23 members.

Figure 2 shows out of these 51 modules, 34 responded in at least one of the antibiotics stresses. The ordering of heatmap from top to bottom separates two regimes that are visible without further analysis and indicated by a black line (Figure 2). Amplitude and breadth are inversely related across the antibiotic arm, so the modules that respond most strongly are also the ones that respond to fewest drugs and are shown at the top of the panel. Fosfomycin, cefuroxime, gentamicin, coumermycin and azithromycin each place at least one module under unique selection to that agent, and fosfomycin and cefuroxime place the largest number of modules under selection overall, 18 each at an absolute activity of 5 or above. Module activity assigned as positive or negative carries distinct mechanistic implications: it identifies members that are required under the stress versus being dispensable.

**Figure 2.**
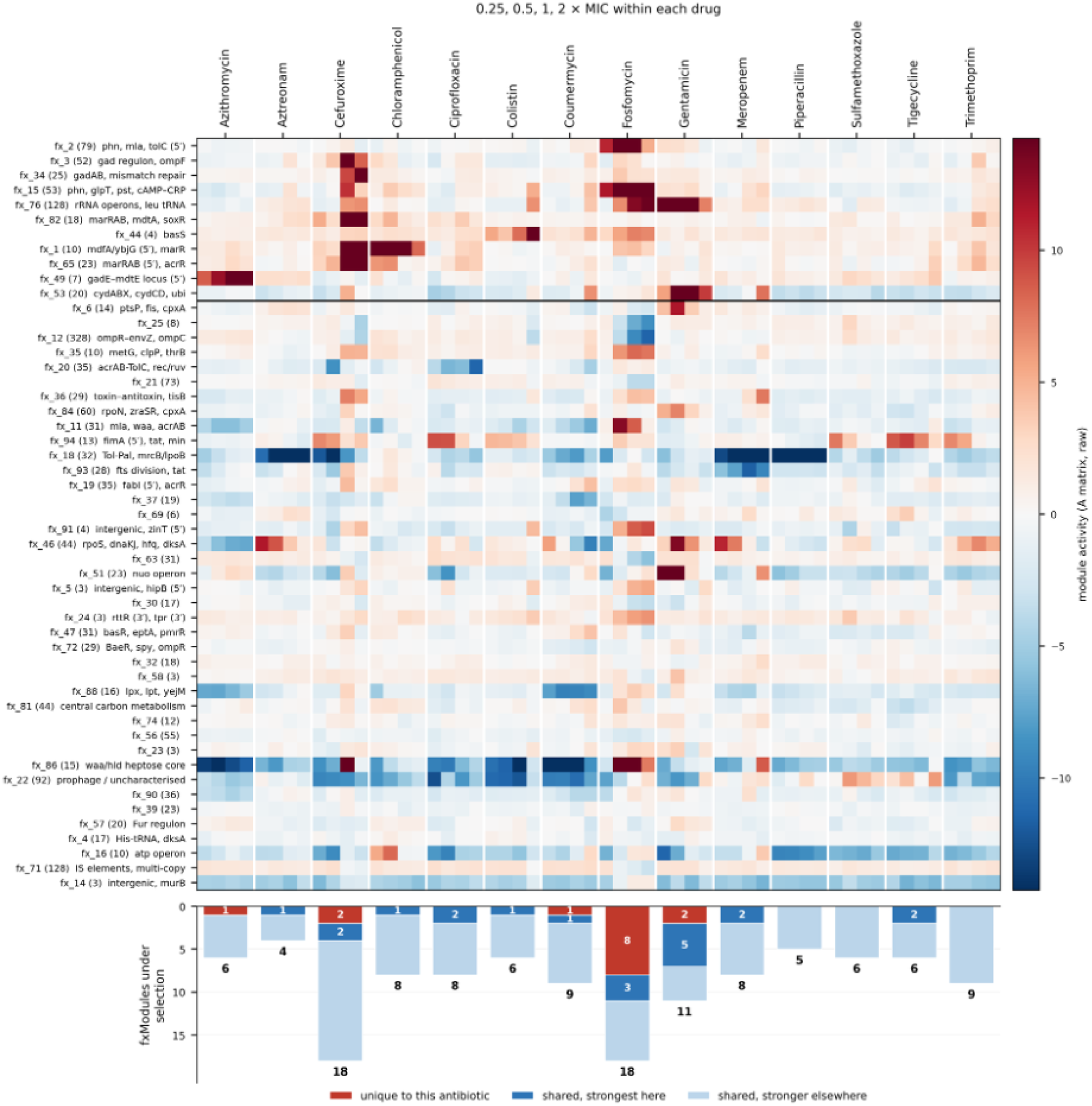
Activity of all 51 retained fxModules across the antibiotic exposures. Columns represent the fourteen drugs, each at 0.25x, 0.5x, 1x and 2x MIC; activities are raw values from the ICA activity matrix and control libraries are not shown. Above the black line are fxModules with antibiotic specific responses and below the black line core modules are shown responding across the panel. In the bottom panel the number of selected modules for each stress are shown; red colour indicates the number of fxModules unique to this stress, dark blue indicates the number of fxModules most strongly associated with this stress, and light blue the number implicated but with a stronger signal for another antibiotic.

Chloramphenicol, colistin, aztreonam, meropenem, ciprofloxacin and tigecycline each dominate selection of at least one module that is unique to them and other drugs may also share the module but lesser intensity. Piperacillin, sulfamethoxazole and trimethoprim act entirely through modules that other antibiotics dominate, trimethoprim crossing the threshold in nine of these modules. Their responses are therefore distributed across shared physiological processes rather than being concentrated in dedicated components relevant to only the selective agent.

### 2.3 Promoters and regulators define chloramphenicol resistance mechanism

One of the unique features of TraDIS-Xpress is its outward facing promoters that can assay an essential gene and indicate overexpression of essential or conditionally important genes. This ICA analysis has identified the contribution of these 5′ and 3′ regions in multiple modules but some of the modules have 5′ and 3′ regions that carry defining weights of the module or the gene bodies where all the insertions are unidirectional indicating overexpression of the neighbouring gene that is essential in the condition but weight is contributed by the overexpressing promoter region. Promoter proximal features make upto 28% of the total feature matrix in the complete database. Ranking the retained modules by the share of total module weight carried by 5′ and 3′ members isolates a small group in which those features dominate rather than merely appear. For example, the strongest activity module of chloramphenicol stress (fxModule_1) was constituted principally by the outward-facing promoter signals. A gene level matrix cannot represent this signal at all, since both modules would collapse onto the flanking gene bodies.

The fxModule_1 contains ten members, with 50% of the module members are promoter members (5′ members), and remaining 5 members have unidirectional insertions (promoter driven insertions) responding to chloramphenicol at different activity levels from 0.25, 0.5, 1 and 2 × MIC with peak activity at 0.5MIC (Figure 3A, 3B). The module’s highest responding members are 5′-*ybjG* and 5′-*mdfA*, but these do not indicate two separate regulatory regions rather one common regulatory control region. As *ybjG* is on the minus strand and *mdfA* on the plus strand, these two 5′ features overlap by approximately 110 bp across the single intergenic control region that separates them (Figure 3A). Therefore, all the reads in *ybjG,* 5′-*ybjG* and 5′-*mdfA* identify only one control region based on all the reads pointing towards *mdfA*. The remaining promoter members are 5′-*acrE,* 5′-*marC* and 5′-*marR.* The 5′-*acrE* represents over expression of *acrEF* operon that encodes AcrEF efflux pump while 5′-*marC* and 5′-*marR* have same intergenic region, 185 bp over lapping, so all the reads are pointing towards *marR*. Moreover, *marR* is part of this module indicating its deletion is beneficial for survival under chloramphenicol stress, so all the reads are unidirectional from *marC,* 5′-*marC* and 5′-*marR* and *marR* indicating over expression of *marA*. The remaining members are the gene bodies of two repressors (regulators) *soxR* and *acrR* (Figure 3A, 3B). SoxR has unidirectional reads that disrupt *soxR* as well as over expressing *soxS* and similarly in *acrR*, all the reads are unidirectional, knocking out *acrR* as well as over expressing *acrAB* encoding *acrAB* pump. MdfA is a chromosomal multidrug transporter of *E. coli* and AcrAB and AcrEF pumps are known for chloramphenicol resistance (Figure 3C). This observation that efflux pump activity can result in decreased chloramphenicol accumulation is well understood (Langevin, El Meouche, and Dunlop 2020) but this unique module highlights the potential of data analysis based on just expression impacts that has not been reported. In principle, this module clearly contains a series of genomically separated genes which are linked by their ability to decrease chloramphenicol accumulation when overexpressed and regulators that control these expressions (Figure 3A).

**Figure 3.**
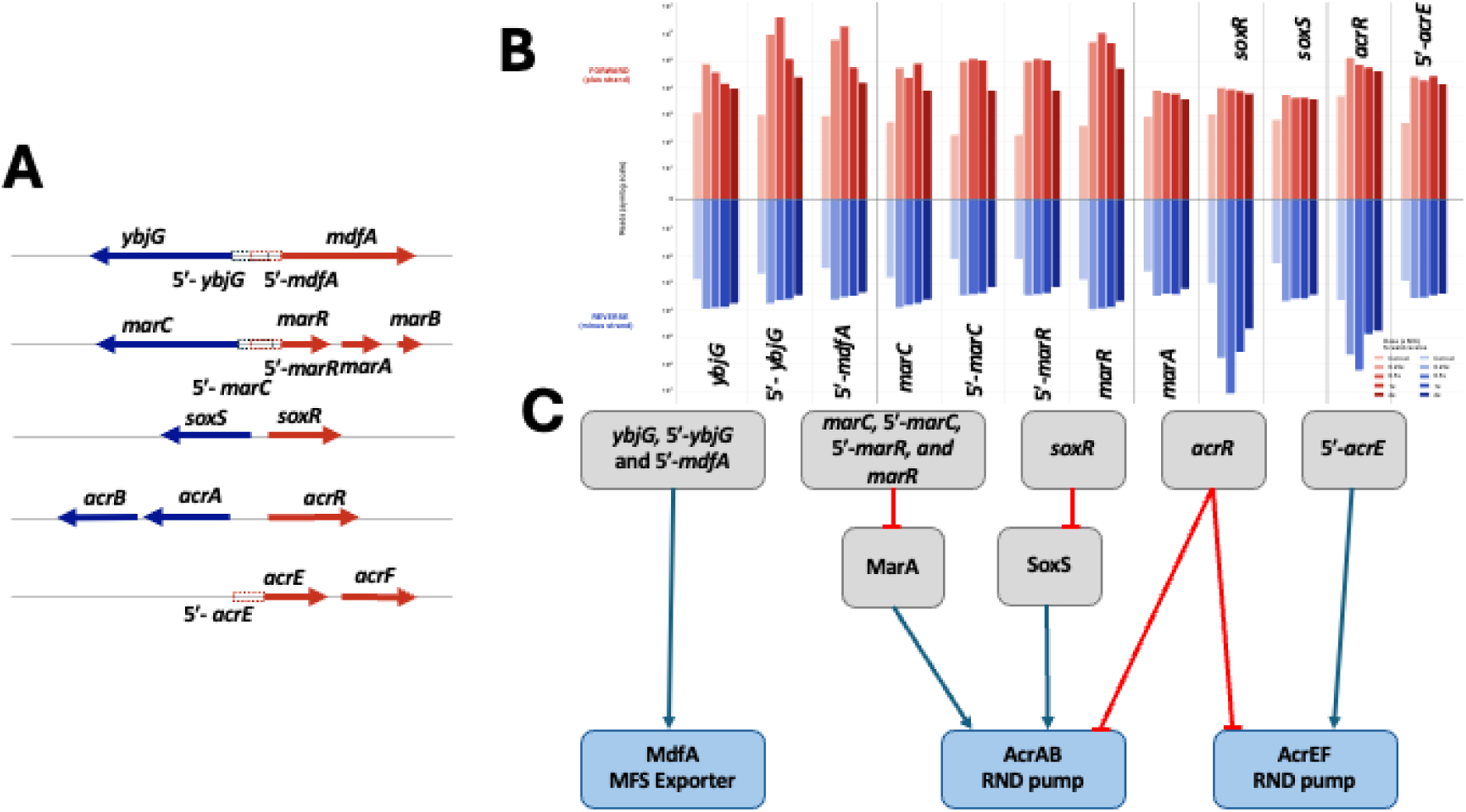
fxModule_1 and the chloramphenicol response. **A.** Position of module members relative to the intergenic midpoint of each divergent or silenced efflux control region **B.** Strand-resolved insertion read distributions for each module feature across chloramphenicol concentrations. **C.** Regulatory model linking module features to the MdfA, AcrAB and AcrEF efflux systems.

### 2.4 Beta-lactam class specific module

fxModule_18 comprised 32 members related to cell envelope responding to all beta-lactam antibiotics, aztreonam, meropenem, piperacillin and cefuroxime (Figure 4A). Instead of corresponding to a metabolic pathway or a regulon, its membership maps almost entirely onto the machinery that synthesises, cleaves and builds the peptidoglycan and structural proteins that span across the cell envelope including the outer membrane. Four sets of physically interacting proteins reported in the literature are recovered as essentially intact modules of the relevant encoding genes. The first is the synthase control complex that spans across the cell envelop, and it includes the complete *tol-pal* operon (*tolQ, tolR, tolA, tolB, pal*) together with *cpoB, lpoB* and *mrcB*, whose products form a single regulatory circuit in which LpoB stimulates MrcB (PBP1B transpeptidase) activity from the outer membrane while CpoB modulates that activity according to the proton motive force transduced by TolQR and TolA (Typas et al. 2010; Gray et al. 2015). The second function is the proteolytic activity controlling hydrolase abundance, comprising *nlpI, prc* and *mepS*, in which the NlpI adaptor both recruits MepS to sites of synthesis and presents it to Prc for degradation (Singh et al. 2015; Su et al. 2017). The third is the Rod complex (*mreBCD, rodZ, mrdA*) that helps in maintaining the rod shape of *E. coli* and its elongation. Moreover, the *mreBCD* operon is essential so it does not tolerate insertions that is why 5′-*mreB* is a member and its upstream genes 3′-*csrD* that have insertions unidirectional over expressing *mreBCD* are observed. The fourth is the septal amidase activation represented by envC.

**Figure 4.**
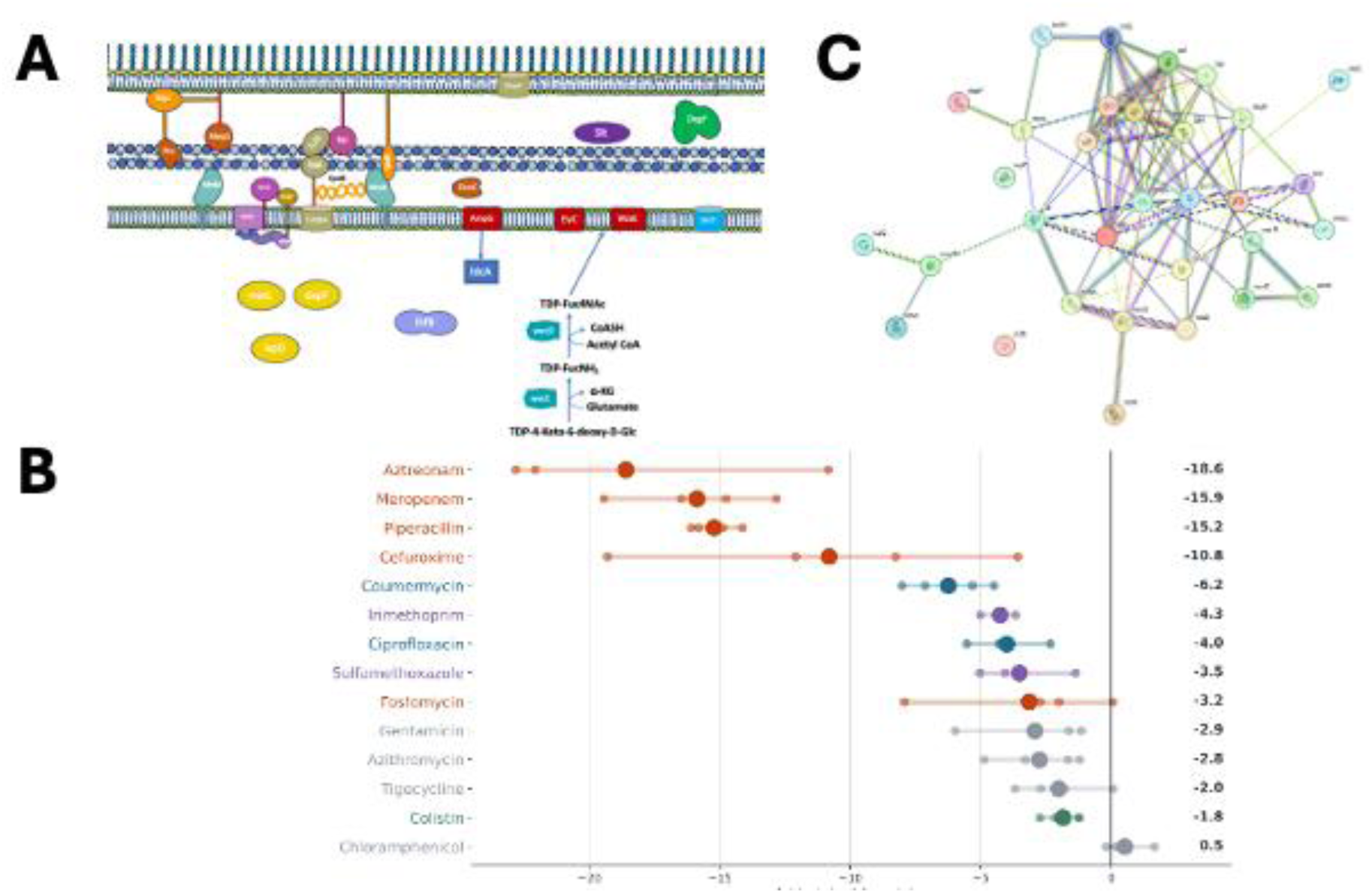
fxModule_18 members. (A) Module members highlighted in relation to their position in cell envelope (B) Module activity across all drugs concentration conditions. (C) STRING DB interaction network of module members.

These physically interacting proteins are supported by two metabolic supply chains that converge on lipid II: a diaminopimelate arm (*dapF,* with *metL* upstream) providing the meso-DAP residue that forms the cross-link targeted by β-lactams, and (*slt, ldcA, ampG*) constitute the muropeptide recycling loop. The undecaprenyl phosphate arm (*ispD, wecE, wecD, wzxE, ycbC*) reflecting competition for the shared lipid carrier used by peptidoglycan and enterobacterial common antigen (ECA) synthesis (Jorgenson et al. 2016; Paradis-Bleau et al. 2014). The *wecE* and *wecD* are important for ECA but mutations in these genes can clog the supply of undecaprenyl phosphate supply. The remaining members, *lpp, degP, secD* and *ompC* contribute envelope tethering, periplasmic proteostasis, export and drug influx respectively.

The module is a typical example as it contains six structural proteins, six transporters, 13 enzymes, four non-catalytic regulators, two proteins of unassigned function (*yajG, ycbc* (ElyC)), and a single transcription factor (*ihfB*). The scarcity of transcriptional regulators is the salient feature. Regulation within this module is through lipoprotein adaptors and activators (*lpoB, ybgF, nlpI, envC*), through targeted proteolysis (*prc, degP*), and through the energetic state of the inner membrane. *ompC* is the only member whose fitness effect is expected to be opposite in sign to the remainder, since loss of the porin should reduce drug influx.

Out of the 32 members of fxModule_18, that are spread across 17 KEGG pathways with only two of the pathways are enriched by FDR less than 0.05. Although the STRING DB analysis shows known or proposed interactions of 30 of the members; with a majority already being linked to beta-lactam resistance. Two of the genes in the module, *ifhB* and *ispD* have unique positions in this network and have not been linked in the STRING DB network.

### 2.5 Fosfomycin drug entry and metabolism

We used the fosfomycin data set for a deeper analysis of the accuracy of predicted fxModules as it has a well characterised mechanism of action and intracellular accumulation. This therefore made a good test of whether module structure reproduces known mechanisms at a resolution that gene set analysis does not reach and if distinct mechanisms can be accurately recalled. Fosfomycin stress induced a very distinct response with more than 18 modules contributing to fitness upon exposure with 8 being unique to this stress. The strongest modules were fxModules_15, 2 and 86 (Figure 5A). fxModule_15 is the highest activity module of the fosfomycin, comprising 53 members. Its membership is dominated by genes that influence the route through which the drug enters the cell.

**Figure 5.**
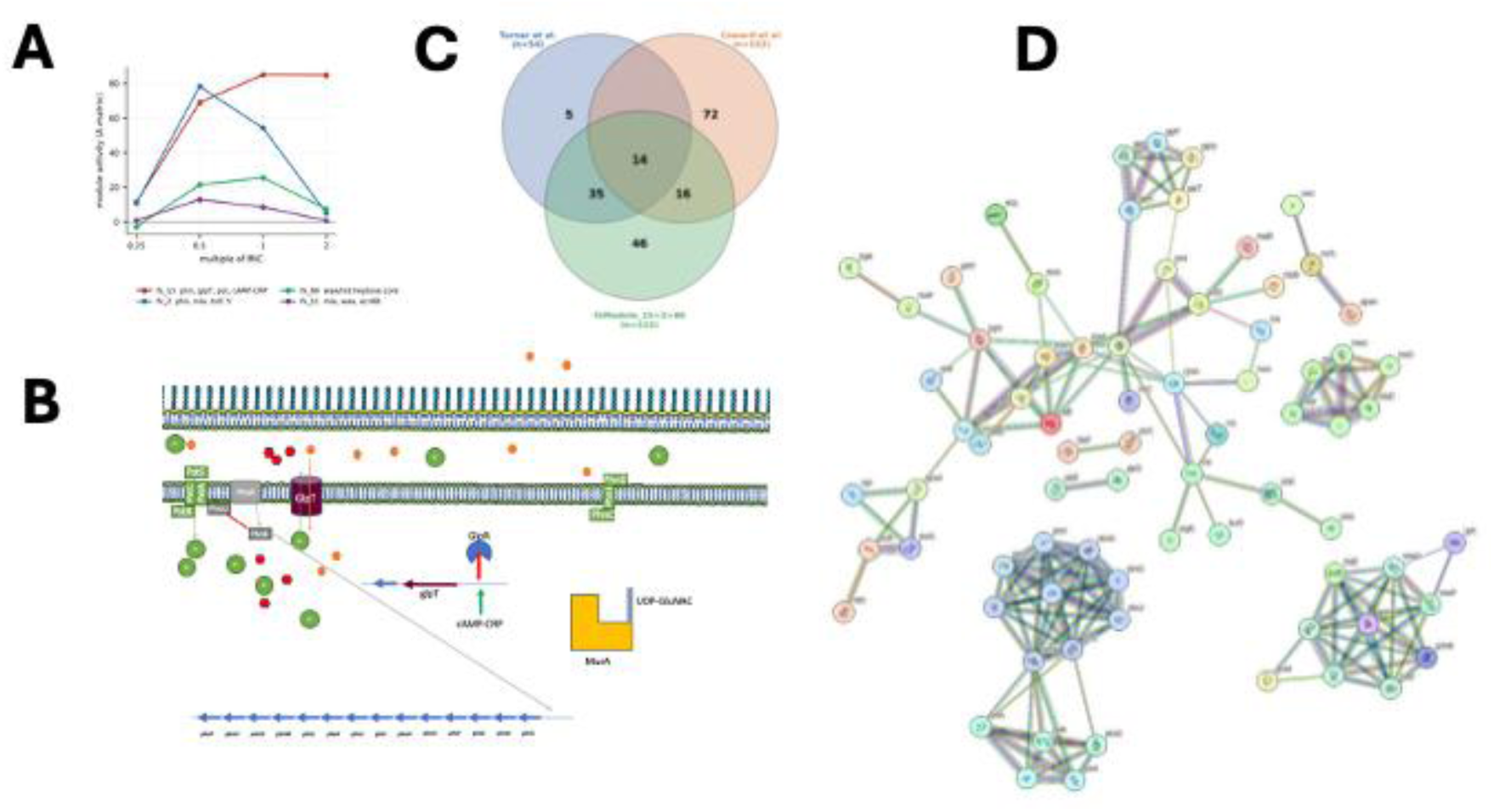
Fosfomycin response. (A) Activity of fxModule_15, fxModule_2, fxModule_86 and fxModule_11 across multiples of MIC. (B) The panel depicts members of fxModules_15 important in transportation and survival of fosfomycin. (C) Overlap of fxModule_15, 2 and 86 members with genes reported by Turner, Yasir et al. (2020) and Coward et al. (2020) (D) STRING interaction network of module members.

In *E. coli* GlpT is the sn-glycerol-3-phosphate (G3P)/phosphate antiporter that also happens to be the principal fosfomycin importer (Figure 5B). This module recovers the *glp* regulon that sets its expression, GlpR that represses the expression and cAMP-CRP that activates its expression. G3P serves as the GlpT co-substrate and as the inducer relieving its transcriptional repressor, GlpR repression of *glpT*. Another enzyme, the GlpD converts this G3P to dihydroxyacetone phosphate (DHAP) that cannot release repressor GlpR. So, for keeping this importer shut, either *glpT* should be knocked out or the repressor should be in excess amount, alternatively, its repressor’s reliever (G3P) should be absent or in the minimum quantity. This module shows knockout of *glpT*, over expression of *glpR* (unidirectional insertions in *glpG*) and over expression of *glpD* (unidirectional insertions in 5’ *glpD*, 5’ *glpE* and *glpE*) keeping the *glpT* down, as well as provide ample amount of repressor GlpR to stop transcription and provide the converter (GlpD) that keeps repressor bound to promoter of *glpT* leading to stop entry in the cell. This is not the only mechanism adopted by cells, it keeps the cAMP–CRP down that activates *glpT* transcription. The *cyaA*, a sole source of cAMP; *crp* and the phosphotransferase relay that activates adenylate cyclase, *ptsI, ptsH*. The *crr* thereby captures the classical indirect resistance route in which inactivation of *cyaA, crp* or *ptsI* silences both *glpT* and *uhpT* without mutating either transporter.

The remainder of the modules comprise paths that both compete with and deliver the drug, converging on the target. The phosphonate operon is present in full, the PhnCDE importer (*phnC, phnD, phnE*) with its regulator *phnF*. The carbon phosphorus lyase complex (*phnG, phnH, phnI, phnJ, phnK*), alongside the complete high affinity phosphate importer and its negative regulator (*pstS, pstC, pstA, pstB, phoU*). Because loss of Pst or PhoU renders the Pho regulon constitutive, activating *phn* while altering the internal free phosphate pool that GlpT exchanges outward.

In *E. coli*,the fosfomycin target is MurA (UDP-N-acetylglucosamine enolpyruvyl transferase) and it inactives MurA by covalently binding at Cys115 as it is a structural analogue of phosphoenolpyruvate (PEP). fxModule_15 contains a cluster of sugar phosphate and purine/nucleotide genes whose connection to fosfomycin runs through the substrate requirements of MurA itself. Fosfomycin does not inactivate free MurA: alkylation of the active site Cys115 requires prior binding of UDP-GlcNAc, which drives the induced fit loop closure that renders the thiol nucleophilic, so fosfomycin binds to MurA-UDP-GlcNA instead of MurA alone. The size of the UDP-GlcNAc pool therefore sets how much of the target is presented to the drug in an attackable state, and that pool is built from two converging precursors. The module’s sugar phosphate members (*treC, galU*) and its purine/nucleotide members (*cmk, guaA, purA, purB*) sit on the supply lines feeding these two precursors. A novel mechanism is through *als* operon, in LB the relevant flux is most probably pentose derived instead of allose, entering via nucleoside catabolism to ribose-5-phosphate and PRPP. Disruptions in the metabolic pathways, divert this flux into phosphorylated dead ends most clearly the derepressed *alsABCEK* branch by knockout of *alsR* with unidirectional insertions. More specifically, the AlsK uses ATP to phosphorylate the sugar and the free phosphate in metabolite are ultimately used to replenish ATP leading to the dearth of pi for GlpT antiporter. There are two steps that block these, allose-6-P is converted to allulose-6-P, by RpiB and allulose-6-P to fructose-6-phosphate is carried out by AlsE. There is a significant knockdown of *alsE* and 5′ *rpiB* indicating the step of conversion is blocked therefore leading to the starvation of phosphates. Another route of action is decrease in Pi pool looks like phosphate starvation to PhoR, which activates PhoB and switches on the *phn* operon, the lyase operon described above further breakdown of fosfomycin as it is a carbon phosphate compound. LeuO regulator overexpression is identified in this module, so Pi starvation might be activating ppGpp and is regulating the module through LeuO although the binding sites on promoters of this operon are not very clearly studied.

The features of fxModule_15 are distributed across several unrelated KEGG assignments. When we compare the analysis here with two recent detailed TIS papers analysing fosfomycin responsive genes in *E. coli* BW25113, a majority of genes identified in those studies (Coward et al. 2020; Turner, Yasir, et al. 2020) are recovered here (Figure 5C). The STRING analysis of fxModule_15 also shows multiple independent clusters that modular analysis has found in the ICA (Figure 5D). Our approach here has accurately identified a module of genes which impact fosfomycin susceptibility and linked genes involved in diverse pathways without the need for extensive manual analysis and data curation.

### 2.6 Respiratory chain components response under gentamicin

Gentamicin is the strongest contributor to fxModule_76 (128 features, activity 83.6) and it comprises the ribosomal RNA operons and leucine tRNAs. As gentamicin has a known target in the ribosome, we will not discuss the details here. Although, respiratory components analysis highlights functional linkage where these pathways intersect to provide benefits in the presence of gentamicin and recalls the particular members of these pathways involved in response to this stress (Figure 6A). The ICA analysis results in three distinct respiratory complexes: fxModule_53 (cytochrome bd-I with ubiquinone biosynthesis, *hemC*, *hemD* and *arcA*; activity 34.1), fxModule_51 (the complete *nuo*operon with *sucA*, *sdhA*, *gltA* and *mdh*; 17.4) and fxModule_16 (the *atp* operon; 11.2), all these components are selected in response to gentamicin (Figure 6B).

**Figure 6.**
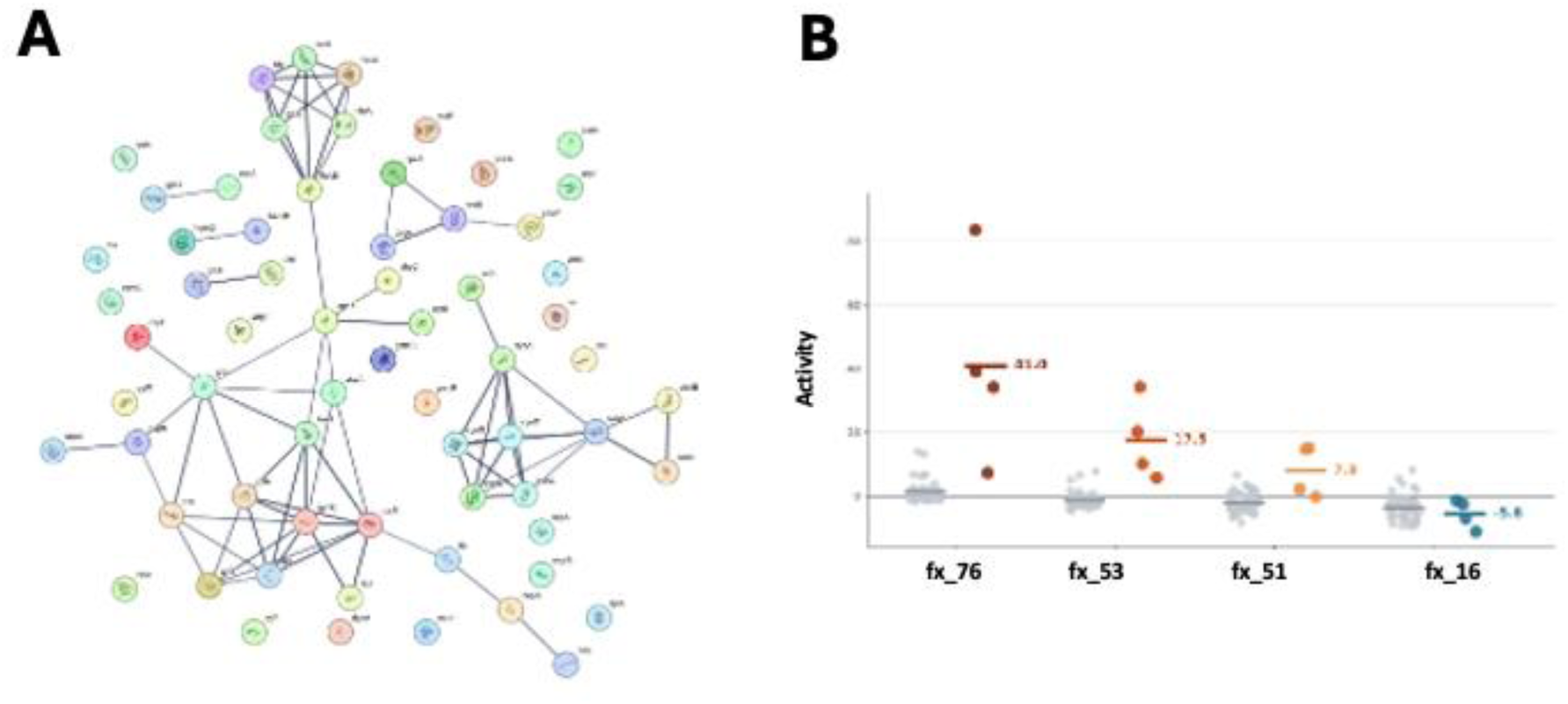
Gentamicin response. **A** STRING interaction network of fxModule_76 members **B** Activity of fxModule_76, fxModule_53, fxModule_51 and fxModule_16 across multiples of MIC of gentamicin in different colours and grey indicate aggregate in other antibiotics.

All the modules appearing related to oxidative phosphorylation seem similar, but the decomposition analysis shows that its components are not interchangeable under gentamicin stress. As PMF is a known mechanism of gentamicin entry from periplasm into the cell. The quinol oxidase and quinone biosynthesis module (fxModule_53) that builds up the PMF across inner membrane has activity twice as strongly to gentamicin stress as the NADH dehydrogenase module (fxModule_51) that provide the proton donors. That *hemC* and *hemD* fall in the same module as the *cyd* and *ubi* genes is consistent with the haem and quinone branches having been identified together in classical aminoglycoside resistance screens. More importantly these activities are positive, indicating the deletion of these module members results in less PMF and thus the gentamicin entry into the cell. On the other hand, other members of oxidative phosphorylation pathway have an independent module (fxModule_16) that include all *atp* genes and its protection increases the survival of the cells under gentamicin stress, as these use PMF to convert ADP to ATP and this reduces the PMF across the inner membrane and gentamicin entry into the cell. One unique observation is the presence of regulators in fxModule_53 and fxModule_51, ArcA and HNS respectively. ArcA has been reported to be involved in regulation of *cyoABCDE* and *cydAB* so its deletion makes cells more susceptible to gentamicin and *hns* deletion has been reported the same, its deletion decreases the survival under gentamicin stress. On the contrary, our data shows *arcA* and *hns* deletion increases the survival chances under gentamicin stress (Chen et al. 2025; Choe et al. 2025).

The four modules do not share a single dose profile (Figure 6B). Activity of the ribosomal RNA module is maximal at 0.5× MIC and that of the *nuo* module at 0.25× MIC, whereas the cytochrome bd-I and ubiquinone module rises steadily to a maximum at 1× MIC. The *atp* module is the only one of the four whose activity is negative under gentamicin at every concentration, and it is closest to zero at inhibitory concentrations. All four converge towards zero at 2× MIC, defining a window between 0.25× and 1× MIC in which the surviving population is still large enough to sample.

## 3. Discussion

Two questions must be answered separately before this analysis can be judged. The first is whether independent component analysis applied to TraDIS-Xpress fitness and expression data gives a stable, reproducible decomposition, and whether the components it returns are real. The second is what biological insight a module level analysis of a TIS compendium provides that a gene or pathway analysis of the same data cannot, and how that insight can be applied.

Taking the first question, the number of components is fixed by ICA dimensionality and convergence tolerance rather than by any downstream choice. Dimensionality was swept from 100 to 550 and set at 400 (Supplementary FigureS1), the point at which the number of robust components across the compendium was maximal, and components were retained only if they recurred across 100 FastICA runs under DBSCAN clustering. Of the solutions obtained at convergence tolerances of 1 × 10⁻⁵, 1 × 10⁻⁶ and 1 × 10⁻⁷, the latter two were identical by direct comparison of source matrices, and the 1 × 10⁻⁷ solution was used throughout in this manuscript. The decomposition we interpret is therefore converged and reproducible rather than a single favourable run.

The membership threshold was calculated using the D’Agostino K² procedure as it was used in the expression (iModulon) and RB-TnSeq fitness (fModule) applications (Borchert et al. 2024; Sastry et al. 2019), and for fitness and expression data it does not change how many modules exist, only which members pass the threshold. We therefore report the sweep from D = 450 to D = 1,850 in full and fix D = 1,350 for all activity analyses, but this choice is empirical and comparison against modules defined at another cutoff must account for it (Supplementary Figure S3). As the cutoff is empirical, it does not leave the pipeline unanchored, a run at D = 1,850 reproduces the fModules described by Borchert et al. (2024), accurately recovering a published decomposition.

The decomposition also behaves internally as expected. At D = 1,350, 51 of the 99 retained modules hold three or more members, with a median of 23, and 34 modules respond in at least one antibiotic exposure at an absolute activity cutoff of 5 or above. Amplitude and breadth are inversely related across the antibiotic arm, the modules responding most strongly being those that respond to fewest drugs, as would be expected if the decomposition separates agent specific selection from shared physiology. Activity also tracks concentration in a component specific way, the four gentamicin responsive modules each peaking at a different multiple of MIC and all converging towards zero at 2× MIC, where replicate agreement falls because the surviving population samples the library sparsely.

The decisive test, however, is whether known mechanisms are recovered from co-fitness and covariation alone, since no annotation, pathway membership or regulon definition enters the decomposition at any stage. The antibiotic arm was chosen for this reason, most of these agents having well described modes of action and thirteen having recently been analysed conventionally (Bastkowski et al. 2026). For fosfomycin, fxModule_15 assembles the route by which the drug enters the cell together with the regulatory circuits that sets it: *glpT* with the *glp* regulon that controls it, and the cAMP–CRP arm (c*yaA, crp, ptsI, ptsH, crr*) constituting the classical indirect resistance route in which loss of adenylate cyclase or CRP silences *glpT* without mutating the transporter (Castaneda-Garcia, Blazquez, and Rodriguez-Rojas 2013). A majority of the genes reported in the two most detailed recent TIS analyses of fosfomycin in this strain background (Coward et al. 2020; Turner, Yasir, et al. 2020) are recovered using modular analysis.

For gentamicin, the direction of response agrees with published single gene phenotypes for every member group for which experimental data exist. The positive activity of fxModule_51 and fxModule_53, meaning insertions in their members were enriched and loss of function advantageous, matches the enrichment of nuo disruptants and of quinone binding and ubiquinone biosynthesis functions under aminoglycoside selection (Girgis, Hottes, and Tavazoie 2009; Shan et al. 2015), the tolerance conferred by deletion of *nuoM, nuoI, sucA, gltA, mdh or sdhC* (Shiraliyev and Orman 2024), the resistance conferred by *ubiA* deletion (El Khoury et al. 2023) and by CRISPRi repression of *ubiD* and *ubiX* (Choe et al. 2025), and the reduced aminoglycoside transport of haem deficient *E. coli* (Bryan and Van Den Elzen 1977; Komeda et al. 2025). The negative activity of fxModule_16 agrees with the only comparable transposon dataset, in which *atpE* and the *atpABCDEFH* operon are fitness genes under aminoglycoside exposure with log fold changes matching the direction we observe (Wellner et al. 2024). Sign agreement is a stronger test than membership agreement, because it is a prediction that could have failed visibly but did not. Agreement extends to physical organisation: thirty of the 32 members of the β-lactam responsive fxModule_18 carry known or proposed STRING interactions, with four sets of interacting envelope proteins recovered intact.

One disagreement with prior work was clear, ArcA and H-NS, the two transcription regulators are members of fxModule_53 and fxModule_51 respectively, and deletion of either has been reported to increase gentamicin susceptibility, whereas the activities here indicate that loss of either increases survival (Chen et al. 2025; Choe et al. 2025). Whether this reflects the pooled competitive context of a transposon library, the compendium centred baseline, or a genuine difference under the conditions assayed is not resolved by these data.

Addressing the second question, the value of the module level view lies in the units it defines. Functional units are assembled across database boundaries: fxModule_53 brings together cytochrome bd-I, the ubiquinone and tetrapyrrole branches supplying its substrate and cofactors and the regulator that activates it, spanning at least five KEGG assignments, while the 32 members of fxModule_18 distribute across 17 KEGG pathways of which only two are enriched at FDR < 0.05. Enrichment against one category returns a fragment of the unit; enrichment against all of them returns statistics with no indication that they describe one thing. Moreover, under gentamicin, oxidative phosphorylation gene set splits it into three modules that do not behave alike, fxModule_53 responding twice as strongly as fxModule_51 and fxModule_16 in the opposite direction, separating the components that generate the proton motive force by which the drug enters the cell from those that consume it.

Resolving expression effects separately from disruption effects is the purpose of the TraDIS-Xpress design and carrying that separation into the decomposition rather than discarding it at the counting stage is what distinguishes fxModules from fitness (fModules) or expression modules (Borchert et al. 2024; Sastry et al. 2019). Promoter proximal features constitute 28% of the feature matrix of the TIS compendium, and between 31% to 43% of retained module content is inaccessible to gene level enrichment due to 5′ and 3′ features. A small group of modules is constituted principally by such features: fxModule_1 and fxModule_49 each resolve intergenic control region into its components rather than assembling genes from across the genome that would have been failed in just gene body component analysis.

Beyond recall of established mechanism, the modules nominate new links for testing, and these arrive with a suggestive mechanism of interaction with the members of the module rather than as unannotated entries near the bottom of a ranked list. For example, fxModule_18, has *ihfB*, occupying a unique position, it is the only regulator in the module and is not connected to the remainder of the module in STRING DB network, it purposes its potential covariation with envelope functions under every β-lactam tested. In fxModule_15, the derepressed *alsABCEK* branch, with knockout of *alsR* and strong knockdown of *alsE* and 5′-rpiB, proposes sequestration of free phosphate in phosphorylated dead ends as a route to reduced GlpT mediated fosfomycin uptake, and overexpression of LeuO in the same module suggests an undescribed regulatory input. Each is a defined hypothesis to test by construction of the individual strain or promoter construct for the measurement of susceptibility or activity and competitive fitness outside the pool.

Because the components were defined across all 754 libraries, which include a range of growth stresses as well as the antibiotic arm analysed here, activities under those conditions can be read against the same definitions. This is the principal practical advantage of decomposing a compendium rather than analysing each experiment separately, and it changes how new data are handled. An experiment processed through the same pipeline is described in coordinates already interpreted, so the interpretive cost of each added condition falls as the compendium grows, inverting the present position in which more conditions produce more separate lists.

In conclusion, the decomposition works in the technical sense that it is reproducible, its parameters are characterised rather than assumed, and it recovers known mechanism and the correct direction of effect without access to annotation. Its value is that it describes an entire TIS compendium in one set of data defined functional and expression units, resolves regulatory signal that gene level analysis cannot represent, and converts a growing collection of independent ranked lists into a common framework in which new conditions and determinants can be placed as they are measured.

## 4. Limitations

The approach requires a TIS compendium of diverse stress conditions, and for fitness and expression modules requires an expression capable transposon design; the remainder of the procedure is general, with the qualification that the membership cutoff must be determined for each data type rather than inherited. Activities are centred on the compendium mean rather than on matched controls, so conditions are compared with one another and not against an absolute baseline.

A variety of stress conditions are required for ICA and it needs computational resources accordingly, however, minimum effective number of data sets and corresponding computational power requirements are outside the scope of this paper and will be discussed in future studies

Module membership is a statement about covariation, not a claim of physical or regulatory interaction; the recovery of known complexes corroborates the method but does not license interpreting every module as a complex. Individual novel assignments, in particular the roles proposed for ihfB in fxModule_18, and LeuO in fxModule_15 require direct genetic validation.

## 5. Methods

### 5.1 Strains, library and growth conditions

All data derive from a single high-density mini-Tn5 transposon mutant library of *Escherichia coli* BW25113 containing approximately 800,000 insertion mutants, with a mean insertion density of one insertion every 6 bp. The transposon carries an outward transcribing, IPTG inducible tac promoter located 3′ of the kanamycin resistance cassette, as described for TraDIS-Xpress (Yasir et al. 2020).

Minimum inhibitory concentrations were determined against BW25113 in broth microdilution in LB media Bastkowski et al. (2026). Antibiotic exposures were performed at 0.25×, 0.5×, 1× and 2× MIC in duplicate for 14 agents except ciprofloxacin and cefuroxime, where one sample failed.

### 5.2 Compendium assembly and sequencing

The compendium comprises 754 TraDIS-Xpress sequencing samples spanning antibiotics, biocides exposure and other growth stress conditions and library controls. Of these, 110 constitute the antibiotic arm analysed here: 13 agents reported previously by Bastkowski et al. (2026) together with chloramphenicol. Cefuroxime and ciprofloxacin are represented by seven libraries rather than eight, one concentration of each having a single replicate.

### 5.3 Read mapping and feature level count matrix

All libraries were processed through a common mapping pipeline for adapter and transposon tag trimming parameters and alignment (Bastkowski et al. 2026). Insertion counts were determined per gene and per annotation across the genome, the gene body, a 5′ region of 198 bp and a 3′ region of 198 bp. Any 5′ or 3′ region overlapping a neighbouring gene by more than 25 bp was removed. The resulting matrix contains 6,311 features: 4,556 gene-body features (4,227 locus tags and 329 features named by base coordinates), 1,219 5′ features and 536 3′ features. To retain the directional information supplied by the outward facing promoter, the counts of 5′ or 3′ regions were split into top (forward) and bottom (reverse) strands and the ratio of 3:7 or 7:3 is used as a filter for directional insertions from the plot files.

### 5.4 Independent component analysis

The feature by sample matrix was decomposed with FastICA, run across the full 754-library compendium rather than on any individual experiment, so that component definitions derive from variation across the whole experimental space. A sweep of dimensionality was set from 100 to 550, and set at 400 after finding the optimum number of robust modules at this dimension across the compendium (Supplementary Figure S1). FastICA was run 100 times, and components were clustered across runs with DBSCAN to retain robust components. Three solutions were generated at convergence tolerances of 1 × 10⁻⁵, 1 × 10⁻⁶ and 1 × 10⁻⁷, yielding 94, 100 and 100 components respectively. The 1 × 10⁻⁶ and 1 × 10⁻⁷ solutions were confirmed identical by direct comparison of source matrices (maximum absolute difference 0.0 across all 6,311 × 100 entries). The decomposition returns a feature weight matrix M and a sample activity matrix A.

### 5.5 Module membership thresholding

Feature membership was assigned by the D’Agostino K² procedure applied to the ICA source weights, as implemented for iModulon definition (Sastry et al. 2019). The cutoff D was swept from 450 to 1,850. All activity analyses use D = 1,350.

One degenerate component per solution, holding approximately 78% of all features at every cutoff (fxModule_9 in the 100-component solution, fxModule_47 in the 94-component solution), was excluded from all downstream analysis. Modules retaining fewer than three features were excluded from Jaccard and composition statistics.

### 5.6 Module activity analysis

Module activity for a sample is the value assigned by the ICA activity matrix A. The activity of a drug concentration is the mean across its replicates. Activities are reported without further transformation. Because ICA centres each component across the full compendium, the zero of the activity scale corresponds to the compendium mean; conditions are therefore compared with one another, and control libraries are not plotted. Modules were ordered in Figure 2 by the effective number of drugs to which they respond. A module was counted as responding to a drug at an absolute activity of 5 or above.

### 5.7 Feature classification and enrichment analysis

Features were classified as inaccessible to gene-level pathway enrichment if they were 5′ or 3′ regulatory features, or gene bodies whose products are annotated as uncharacterised or putative, mobile elements, or structural RNAs. KEGG pathway enrichment was performed on module gene body members and in case of 3’ or 5’ feature, gene name was used instead of the feature. Protein interaction networks were retrieved from STRING. Regulon enrichment used a Fisher exact test with Bonferroni correction using R packages.

## Funding and acknowledgements

The authors gratefully acknowledge the support of the Biotechnology and Biological Sciences Research Council (BBSRC); MY, AKT, SB, AT, ERH, IGC and MAW were supported by the BBSRC Institute Strategic Programme Microbes and Food Safety BB/X011011/1 (and its constituent project BBS/E/F/000PR13635) and by the BBSRC Core Capability Grant BB/CCG2260/1.

## Author Contribution

MY, IGC, and MAW, conceived the idea, MY and MAW wrote the main manuscript text, MY developed the computational method and performed the data analysis, AT advised and developed the part of pipeline to handle large data sets, SB provided QuaTraDIS support and pre-processed data, AKT, IGC, and MAW helped in data interpretation and cross validation. ERH helped in expression fxModule interpretation and validation, all authors reviewed the manuscript.

## Data and code availability

The data can be accessed from Bastkowski et al 2026 repository of EBI under project accession number PRJEB29311.

## Supplementary Figures

**Supplementary Figure S1:**
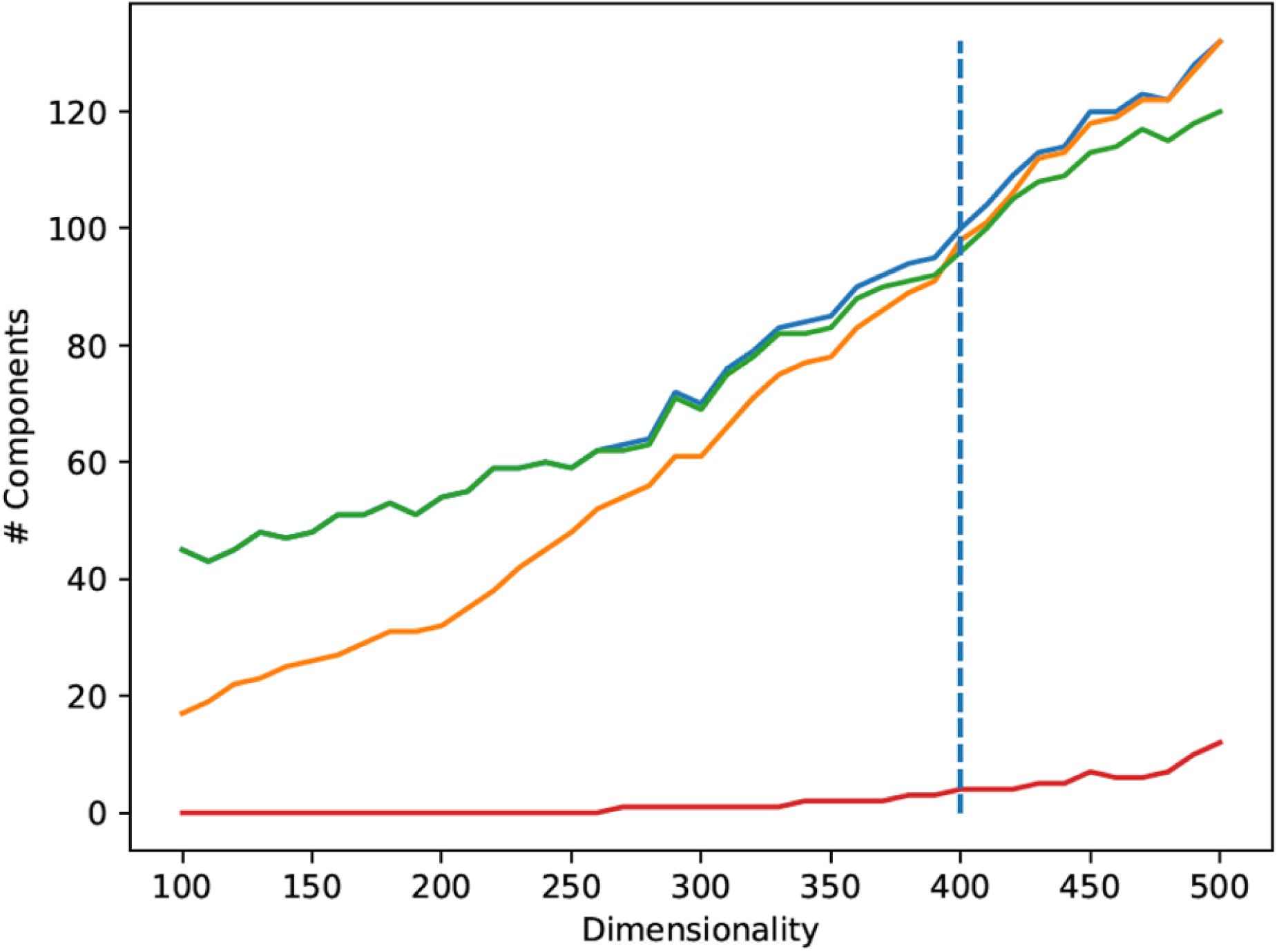
A sweep of fxmodules across dimensions from 100-550. The panel shows the number of robust module peak at 400 and then the single member module start increasing after 400 shown in as red line.

**Supplementary Figure S2.**
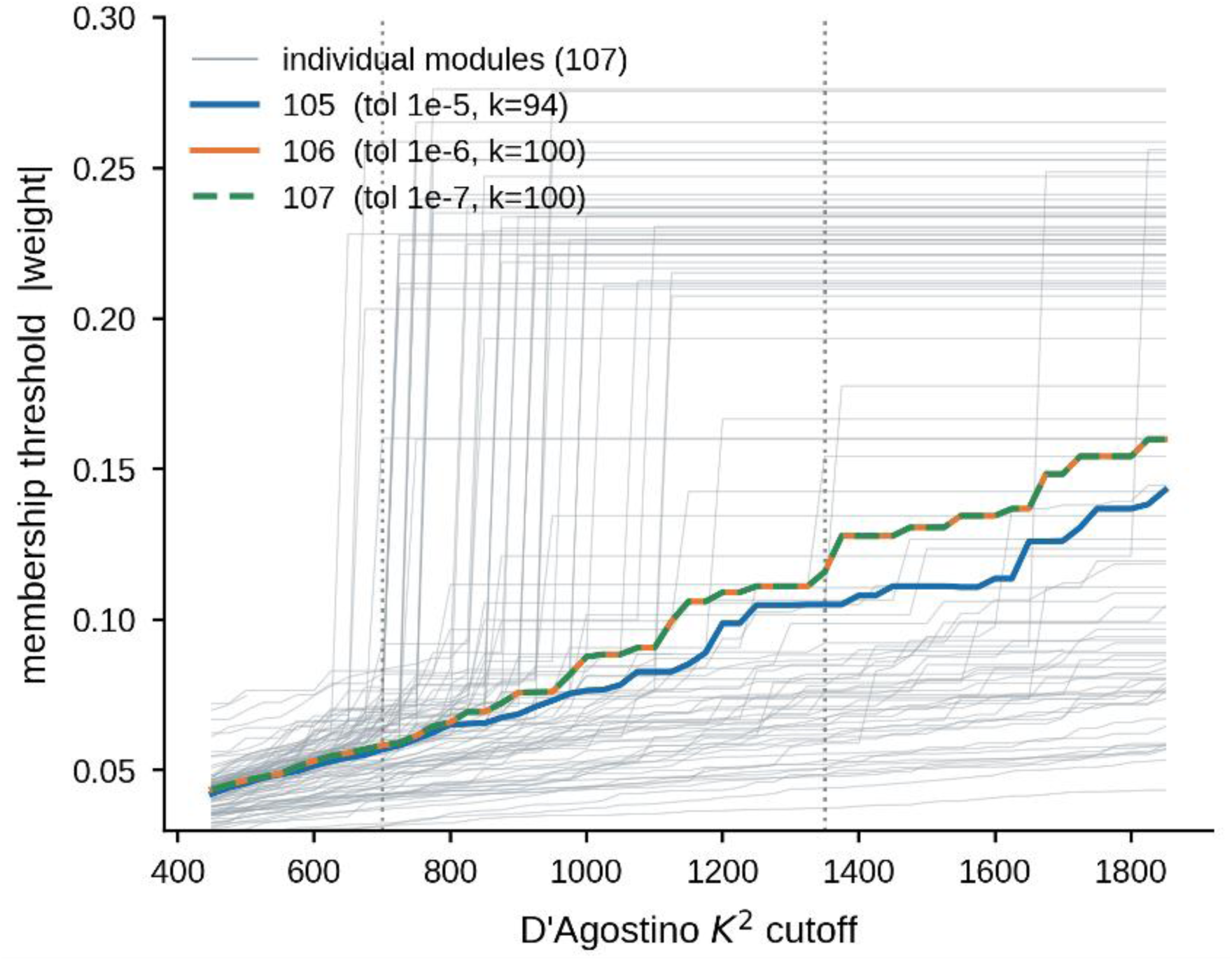
D Agostino K^2^ cutoff for each module: The panel above shows the D Agostino values for each individual module and the optimum number for this data is 1350, shown by vertical dashed line

**Supplementary Figure S3:**
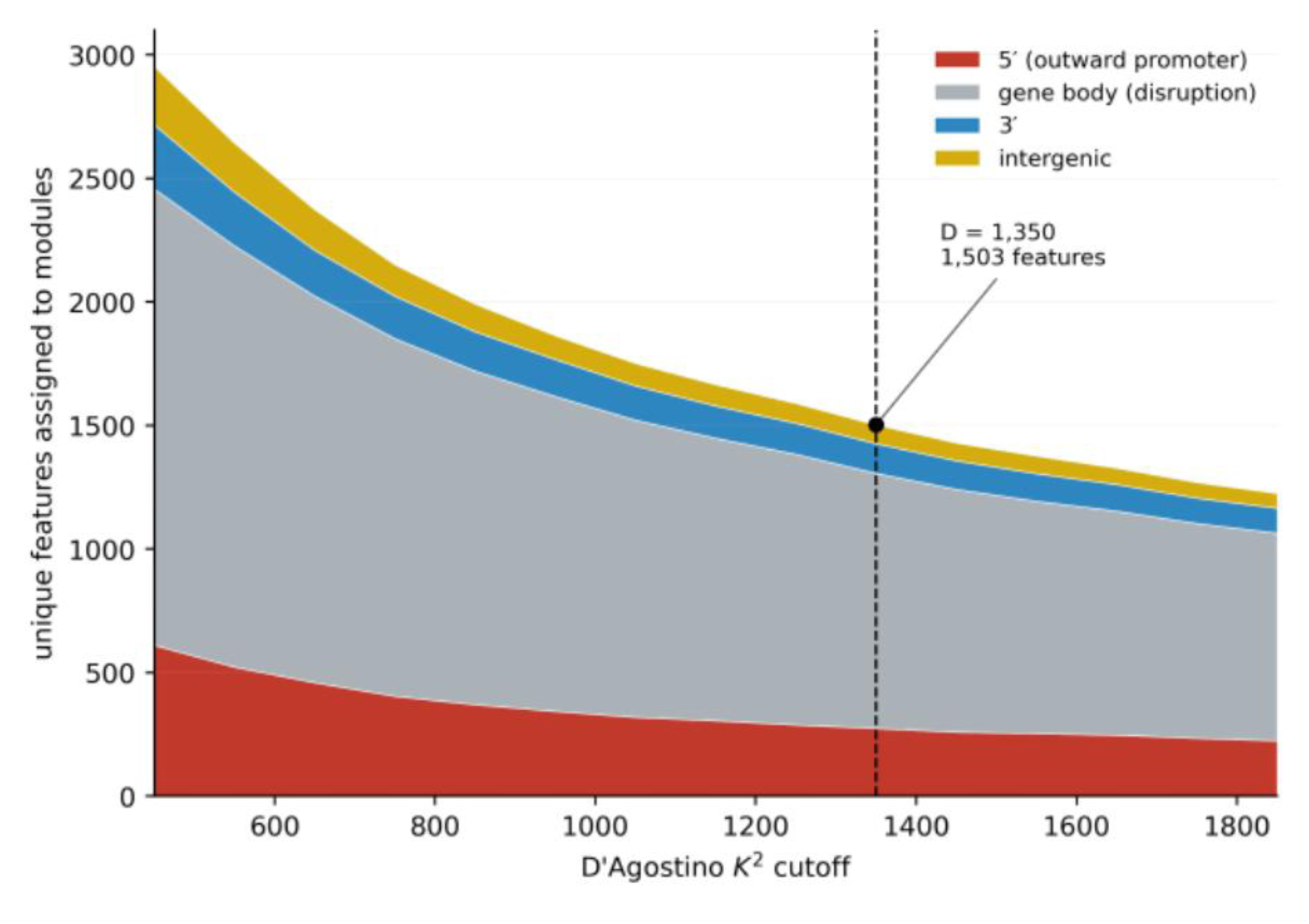
Gene body and 5 and 3 region members of the fxModules: The total members of the modules vary according the to the D Agostino cutoff and D-1350 has resulted in 1503 module members.

